# The changing contributions of weakness and the flexor synergy to post-stroke arm function over time: A kinematic re-examination of Twitchell

**DOI:** 10.64898/2026.02.03.703629

**Authors:** Inbar Avni, Ahmet Arac, Noy Goldhamer, Reut Binyamin-Netser, Shilo Kramer, Simona Bar-Haim, John W. Krakauer, Lior Shmuelof

**Author notes:** Correspondence to: Inbar Avni, Mercy Pavilion, 1622 Locust St, Pittsburgh, PA 15219, USA.

## Abstract

In 1951, the neurologist Thomas Twitchell published a seminal paper in *Brain* describing the time-course of recovery from hemiplegia after stroke in 25 participants from hospitalization to when they reached what he deemed steady state. His main emphasis was on the evolution of voluntary movements at the shoulder, elbow and hand, first within an obligatory flexor synergy, and then independently out-of-synergy. We thought that 75 years later, an update using modern motion capture technology should be attempted as it would allow for finer granularity in the characterization of the time courses of both functional recovery and of the flexor synergy, and then relate them to each other, to weakness and to well-established clinical scales. To this end, we used marker-less 3D kinematics to assess task performance and intrusion of synergies in thirty-three stroke participants longitudinally, from the early sub-acute stage (1 – 8 weeks post-stroke) to the chronic stage (24 – 64 weeks post-stroke). Participants performed an out-of-flexor synergy (shoulder flexion and elbow extension) reaching task. We assessed the time course of recovery of obligatory intrusion of pathological synergies based on measures derived from the angular velocity profiles of the shoulder and the elbow joints. Task-related kinematic measures were obtained and compared to sixteen healthy controls. Grip strength, Motor impairment (FMA), and function (ARAT) scores were also collected. Task kinematics were different from controls in the early, late sub-acute, and chronic stages, but showed gradual recovery over time. Weakness in the hand remained impaired at all time points. Flexor-synergy intrusion was maximal in the early sub-acute stage and then began to subside. Regression analysis with functional kinematic and clinical (FMA, ARAT) measures indicated that flexor-synergy intrusion was a significant predictor in the early and late sub-acute stages, but not in the chronic stage, while weakness remained a significant predictor at all stages of recovery. To better address the relationship between synergies, weakness, and function, we analyzed the more severe cases (ARAT<21) separately. In the sub-acute stage, most of them (11/13) suffered from intrusion of synergies, whereas in the chronic phase, only a minority (2/8) did. Weakness seemed to be the main contributor to poor outcome in the chronic phase. We conclude that weakness and synergy intrusion evolve separately from the subacute to the chronic phase, perhaps more so when neurorehabilitation is given at a dose higher than standard of care.

## Introduction

Stroke is the leading cause of disability in adults.^1^ Upper limb motor impairment after stroke – arm “paresis” - is comprised of multiple components, including weakness, loss of dexterity, spasticity, and the intrusion of pathological synergies.^2^ While longitudinal studies have shown partial recovery of motor function within the first three months post-stroke,^3–5^ how resolution of specific components of arm paresis contributes to this recovery remains poorly understood.

In 1951, Twitchell^6^ proposed that upper limb recovery after stroke follows an ordered sequence: initial improvement in weakness occurs alongside the emergence of pathological synergies, subsequently followed by the resolution of synergies, increased grip strength, and finally, improved dexterity. Despite the continued influence of Twitchell’s findings on our current framing of stroke recovery, a more granular analysis of the time-varying relationship between positive and negative deficits during prehension recovery has not been undertaken up to the present day. In this study, we used 3D marker-less kinematics to track the parallel emergence and resolution of synergy intrusion and weakness, and then related these two time-courses to functional outcomes.

Abnormal synergies, first described by Twitchell as multi-joint obligatory movement patterns occurring with voluntary motion, are considered a positive symptom of stroke.^6^ In the upper limb, voluntary shoulder flexion often leads to involuntary flexion of the elbow and wrist, a pattern referred to as a flexor synergy. Studies have reported conflicting findings regarding the onset and resolution of flexor synergies post-stroke. While some studies suggest an early emergence of synergies,^7,8^ others indicate that synergy emerges later, in the chronic stage.^9–11^ Notably, many of these studies used indirect assays for synergy, such as joint individuation paradigms^7,10,11^ or horizontal plane reaching tasks with anti-gravity or robotic support.^8^ These approaches do not directly assess for synergy intrusion during ongoing execution of a functional task. Recently, we developed such a task, along with a kinematic analysis method, to directly detect flexor synergy intrusion during functional tasks, confirming and quantifying their appearance in the early subacute stage of recovery.^12^

Longitudinal studies on negative signs suggest that spontaneous recovery typically plateaus within the first three months post-stroke. For example, some studies found that motor dexterity, plateaued at five to eight weeks post-stroke.^13,14^ Some studies demonstrated a similar recovery window within the first three months when measuring upper limb weakness,^7,15^ whereas others demonstrated a longer recovery window.^13^ These findings suggest that dexterity and strength recovery is largely complete within the early subacute stage.

In this longitudinal study, we revisited Twitchell’s qualitative observations by investigating how flexor synergies emerge and resolve during recovery and determined their relationship to improvements in grip strength, to standard clinical scales and to kinematic performance in functional tasks.

## Materials and methods

### Participants

The study was conducted at the Adi Negev-Nahalat Eran rehabilitation hospital in collaboration with the Lillian and David E. Feldman Research Center for Rehabilitation Sciences. Research protocols for both stroke and healthy participants were approved by Sheba Hospital Helsinki Committee and Ben-Gurion University Human Participants Research Committee, respectively. All participants were recruited from the inpatient neurological rehabilitation department and provided a written informed consent prior to enrollment in the study. The mean hospitalization period was 53 days.

Data were collected from 33 stroke participants (aged 65.1□± □11.8), starting from the early sub-acute stage to the chronic stage (time points: 2, 5, 12, 26, and 52 weeks). The characteristics of the first time point for each of the participants are presented in Table 1. To control for the missing data points between time points and reduce the number of multiple comparisons, we also used a three-stage approach for assessing group and longitudinal differences: early sub-acute, late sub-acute and chronic (Table 2). This was achieved by taking the earliest recording in the study (early sub-acute) and earliest recording in the chronic stage (> 24 weeks), as well as the 12-week measures for each participant. All participants had data points in the early sub-acute and chronic stages (n = 33). Data from three participants were missing from the late sub-acute stage (n = 30) due to scheduling difficulties. The description of the longitudinal data before and after data aggregation is presented in Table 2.

**Table 1.** Characteristics of participants in the sub-acute phase after stroke.

| Participant number | Age at stroke | Gender | Dominant hand | Paretic side | Stroke type | Weeks post stroke | Spasticity (MAS) | Weakness (kg) | FMA | ARAT |
| --- | --- | --- | --- | --- | --- | --- | --- | --- | --- | --- |
| 1 | 76.0 | Female | R | L | Ischaemic | 1.7 | 1 | 9.0 | 53 | 49 |
| 2 | 68.3 | Female | R | R | Ischaemic | 3.3 | 0 | 15.6 | 65 | 52 |
| 3 | 72.3 | Male | R | R | Ischaemic | 5.1 | 1 | 23.3 | 60 | 46 |
| 4 | 83.7 | Male | R | R | Haemorrhage | 7.3 | 2 | 5.0 | 52 | 49 |
| 5 | 49.1 | Male | R | R | Ischaemic | 5.1 | 2 | 10.0 | 52 | 29 |
| 6 | 66.2 | Male | R | R | Ischaemic | 2.3 | 0 | 32.3 | 65 | 55 |
| 7 | 68.9 | Female | R | R | Ischaemic | 3.7 | 0 | 15.3 | 66 | 49 |
| 8 | 69.8 | Male | R | R | Ischaemic | 7.3 | 0 | 30.0 | 61 | 31 |
| 9 | 50.7 | Male | R | R | Haemorrhage | 8.3 |  | 3.0 | 45 | 36 |
| 10 | 83.7 | Male | R | L | Haemorrhage | 2.1 | 0 | 18.0 | 57 | 38 |
| 11 | 49.1 | Male | L | R | Haemorrhage | 1.4 | 1 | 0 | 0 | 0 |
| 12 | 72.8 | Male | R | L | Haemorrhage | 2.0 | 0 | 20.6 | 58 | 52 |
| 13 | 68.5 | Female | R | L | Haemorrhage | 3.9 | 0 | 0 | 41 | 19 |
| 14 | 64.7 | Male | R | R | Ischaemic | 1.3 | 1 | 0 | 10 | 0 |
| 15 | 44.2 | Male | R | R | Haemorrhage | 3.7 | 2 | 0 | 8 | 0 |
| 16 | 69.4 | Male | R | R | Ischaemic | 2.4 | 0 | 0 | 4 | 0 |
| 17 | 76.0 | Male | R | L | Ischaemic | 3.4 | 0 | 18.4 | 55 | 55 |
| 18 | 56.5 | Female | R | R | Haemorrhage | 3.1 | 0 | 0 | 25 | 0 |
| 19 | 56.1 | Male | R | L | Ischaemic | 2.0 | 0 | 29.3 | 63 | 57 |
| 20 | 64.5 | Male | R | R | Ischaemic | 4.7 | 0 | 33.6 | 64 | 56 |
| 21 | 65.2 | Male | R | L | Ischaemic | 2.1 |  | 32.6 | 63 | 48 |
| 22 | 68.2 | Female | R | L | Ischaemic | 2.1 | 1 | 0 | 35 | 0 |
| 23 | 61.4 | Male | R | L | Ischaemic | 5.4 | 0 | 26.0 | 62 | 56 |
| 24 | 59.8 | Male | R | L | Ischaemic | 3.6 | 1 | 0 | 4 | 0 |
| 25 | 30.1 | Male | R | R | Haemorrhage | 2.0 | 1 | 16.0 | 62 | 55 |
| 26 | 66.1 | Female | R | R | Ischaemic | 3.9 | 0 | 18.7 | 66 | 57 |
| 27 | 81.3 | Male | R | R | Ischaemic | 6.4 | 1 | 0 | 4 | 0 |
| 28 | 53.7 | Male | R | L | Ischaemic | 5.6 | 0 | 20.0 | 58 | 57 |
| 29 | 74.1 | Male | R | L | Ischaemic | 4.3 |  | 0 | 4 | 0 |
| 30 | 77.4 | Female | R | R | Haemorrhage | 3.9 | 0 | 16.6 | 66 | 57 |
| 31 | 74.3 | Male | R | R | Ischaemic | 5.0 | 0 | 36.0 | 63 | 55 |
| 32 | 64.7 | Male | R | R | Haemorrhage | 5.1 | 1 | 1 | 26 | 2 |
| 33 | 60.7 | Male | R | L | Ischaemic | 1.9 | 2 | 1.4 | 36 | 30 |

**Table 2.**
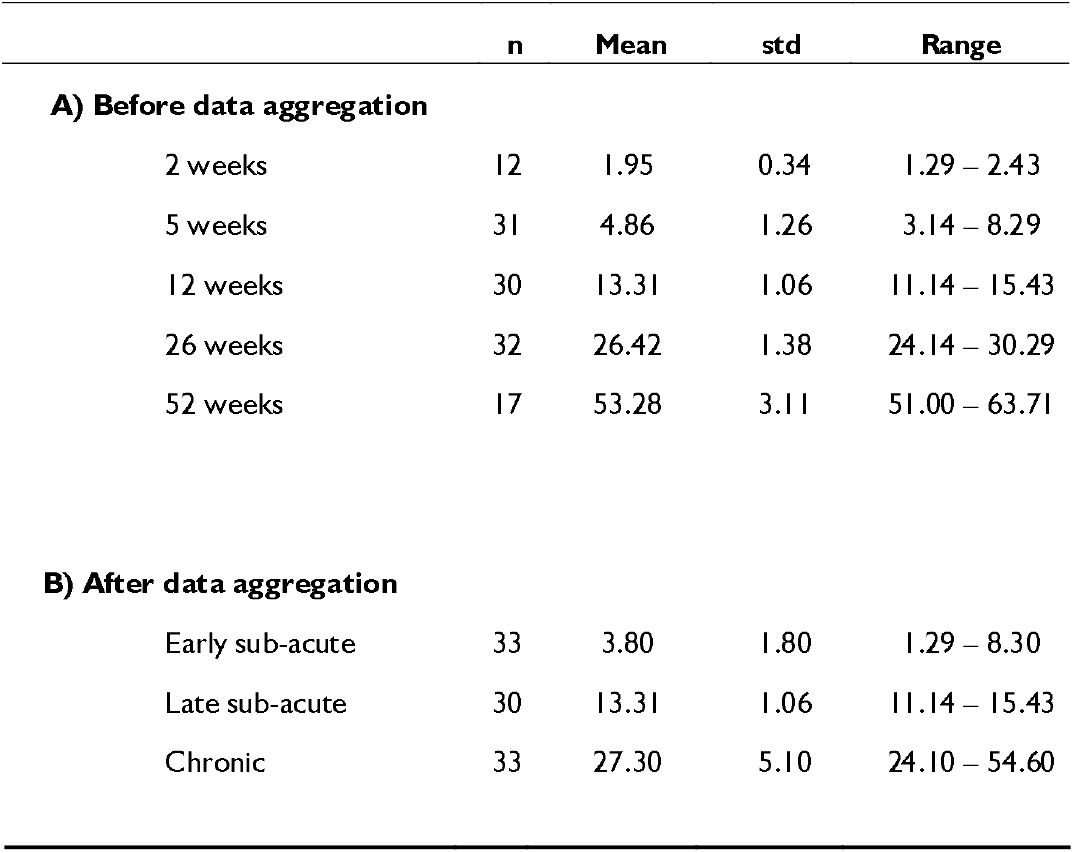
Characteristics of natural history time points.

All participants were able to perform shoulder elevation (as assessed by a trained clinician). The kinematic data of stroke participants were compared to 16 healthy controls (aged 68.8□± □3.5). This study reports longitudinal results of the same group of participants previously examined during their sub-acute stage (n = 33).^12^

### Experimental design

As previously reported,^12^ we recorded participants performing a reaching task with each hand. Participants were instructed to perform upward and forward reaching movements with the hand toward a suspended target. Participants were instructed to complete ten iterations of the task, which required performing a movement pattern of shoulder flexion and elbow extension.

### Recording setup

A two-camera custom-made system was used to record and extract the 3D Kinematic data (DeepBehavior).^16^ This algorithm enabled the detection of joint positions during movements without markers placed on the participants. This marker-less 3D model was based on a convolutional neural network pose estimation algorithm (OpenPose).^17^ The camera system (150 frames per second, 1280×1024 pixels, Blackfly S Color 1.3 MP USB3 camera with a Fujinon 1.5MP 6mm C Mount lens) was positioned 120 cm from the participant at a height of 95 cm with 66 degrees in between the optical axes of the cameras.

Participants were recorded from the front angle while facing the cameras and reaching up towards a suspended object (∼1.5 m above ground). Task recordings were included in the analyses if at least two movements were detected.

### Impairment and functional measures

FMA,^18^ Action Research Arm Test (ARAT),^19^ and strength (grip dynamometer) scores were collected. Strength scores were marked as zero if the arm was flaccid or the participant couldn’t perform a power grip. Grip dynamometry values of each participant were normalized into z-scores according to age, gender, and hand dominance.^20^

## Data analysis

3D kinematic data of 57 body key points were obtained using the DeepBehavior algorithm.^16^ Data were preprocessed using a Savitzky-Golay filter with a window size of 57 and a polynomial degree of 3. Individual movements were detected (local maxima detection algorithm) and segmented based on the wrist tangential velocity profiles (movement start and end were defined based on the crossing point of 10% of the peak velocity). The movement segmentation algorithm was manually verified and adjusted for all participants, and movements.

Several kinematic performance measures were calculated. Extent was computed as the 3D distance between the positions at the end of the reaching task and the start of the movement. Movement duration was defined as the difference between movements’ start and end times in seconds. Peak velocity was defined as the maximum velocity between the start and end in meters per second. Smoothness was defined as the minus log of the normalized integrated squared jerk divided by length^2^/duration^5^ for each movement (see equation 1):

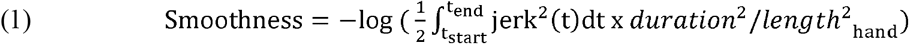

Shoulder and elbow joint angles were calculated using an intrinsic (anatomical) coordinate system (in relation to a specific joint). The shoulder flexion angle is the shoulder angle projected onto the sagittal plane of the subject. To calculate this, we first defined the mid-sagittal plane which is orthogonal to the plane formed by both shoulder and mid-hip key points, and that passes through the mid-hip and mid-chest key points. We then projected the ipsilateral shoulder and elbow key points onto this plane. The change in shoulder flexion angle is then calculated as the angle between the two lines formed by the projected shoulder and elbow key points at the start of the reach and at other time points during the movement. The elbow angle was defined as the angle between the two lines formed by the ipsilateral wrist and elbow, and elbow and shoulder.

As previously reported, two measures of the flexor synergy were quantified based on the angular velocity of the elbow and shoulder joints.^12^ The first measure was the flexor synergy proportion, calculated as the time spent simultaneously flexing the elbow and shoulder divided by the total time of the movement. The second measure was the flexor synergy strength, defined as the Pearson’s correlation coefficient between shoulder flexion and elbow extension angles. This measure was calculated during the longest segment of continuous shoulder flexion found within the movement. The Pearson coefficients were converted using the Fisher transformation (inverse hyperbolic tangent) to deal with this measure’s skewed distribution.

### Statistical analysis

Statistical analysis was performed in two ways. The first was meant to test the level of motor function at different stages of the recovery time course (five time points as denoted in Table 2A). This was achieved by using two-sample t-tests to assess the differences between the kinematic measures recorded in the stroke participants, in each of the five time points, to age matched controls. In addition, a one sample t-test was used to assess the difference of the clinical measures compared to normal motor function (ARAT max score, and dynamometry normal population distributions) recorded in the stroke participants, at each of the five time points (time points as denoted in Table 2A). Significance was adjusted to p=0.01 to correct for the number of comparisons (n = 5).

The second approach was meant to test the recovery time course and was assessed using paired t-tests for detecting group differences between the early sub-acute, late sub-acute, and chronic stages. The differences between groups for each of the three time points (early sub-acute, late sub-acute, and chronic as denoted in Table 2B) was performed using a two-sample t-test with unequal variance, with a power of 82% for an effect size of 0.8. Significance was adjusted to a new p-value to correct for the number of comparisons (p = 0.05/3).

Linear regression analysis was performed to assess the contribution of impairment measures (strength and intrusion of flexor synergy strength) to performance measures (e.g., Fugl-Meyer, ARAT, peak velocity, and extent) in stroke participants. The power for the multiple linear regression analysis was 92% for an effect of f^2^ = 0.3 with two predictors in the model. Statistical analyses were performed using MATLAB^21^ and JASP,^22^ and statistical power was computed using G*power version 3.1.9.4.^23^

## Results

### Flexor synergy intrusion was more of a factor in the early and late sub-acute stages compared to the chronic stage

First, we examined flexor synergy intrusion from the early sub-acute stage to the chronic stage. The intrusion of flexor synergy was quantified using two measures we previously used in a cross-sectional study in the sub-acute stage.^12^ We first assessed the duration of the intrusion by calculating the proportion of movement time during which the elbow and shoulder joints were simultaneously flexed. Secondly, we quantified the strength of the intrusion by computing the Pearson correlation between elbow and shoulder flexion angles during the longest segment of continuous shoulder flexion. Both measures were calculated based on the paretic sides of stroke participants.

The time course of these measures is presented in Figure 1. Both measures were abnormal on the paretic side of stroke participants compared to controls in the sub-acute stage, indicating presence of worsened flexor synergy in stroke participants. The proportion of shoulder flexion-elbow flexion during the reaching movement was higher in stroke compared to controls at the early and late sub-acute stages (*P* < 0.01). In addition, flexor synergy strength was higher in controls in the early sub-acute stage (*P* < 0.001). The same comparisons (for both measures) in the chronic stage were not significant (Figure 1).

**Figure 1.**
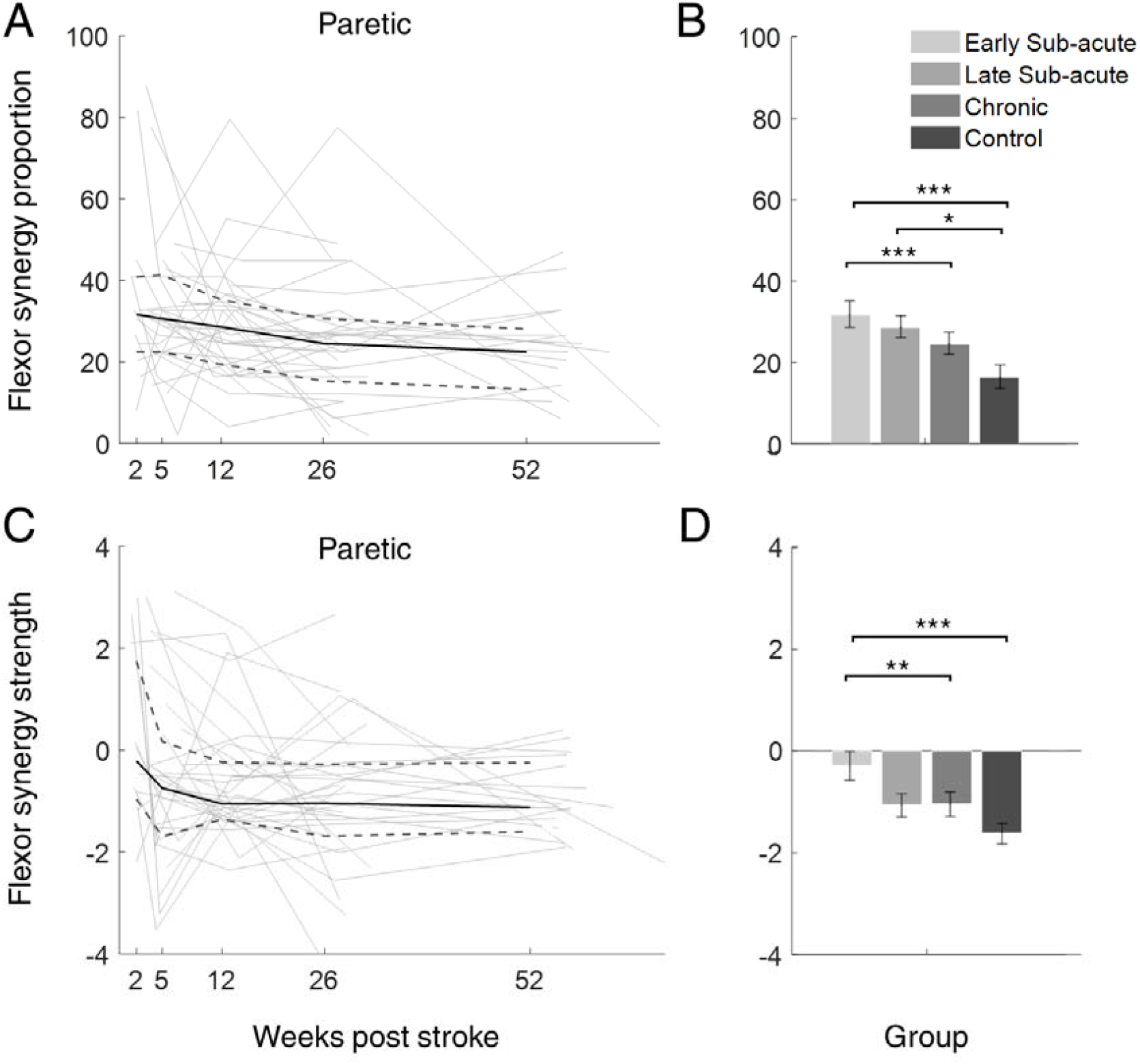
Synergy measures of the paretic and non-paretic arms in a reaching task. **(A-B**) Flexor synergy proportion time course plots and bar plots. (**C-D)** Flexor synergy strength time course plots and bar plots. Left column: Time course plots of the synergy measures of the paretic arm recorded at different stages of recovery. Each line connects the dots of the values of the measure for a single participant across all time points. Group median values are represented by the bold line, and the group 25^th^ and 75^th^ percentile values are represented by the bottom and top dashed lines, respectively. Right column: Group median values of all participants in the early sub-acute stage (1-8 weeks), late sub-acute stage (11-15 weeks), chronic stage (24-56 weeks), and age-matched controls are denoted in the bar plots. Significant group differences are denoted by asterisks (* = *P* < 0.0125, ** = *P* < 0.01, *** = *P* < 0.001, two-sample t-test with unequal variance).

Recovery from flexor synergies was also apparent in significant differences between the intrusion measures in the early sub-acute and chronic stages (*P* < 0.01) (Figure 1 right panel). Thus, intrusion of flexor synergies emerges from the early sub-acute stage and cannot be seen at the chronic stage.

### Abnormal kinematics during task performance persisted in the sub-acute and chronic stages

To quantify the progression of functional impairment after stroke, we extracted several kinematic measures, including movement duration, peak velocity, movement extent, and smoothness (Figure 2). All kinematic measures were abnormal in the sub-acute and chronic stages, except movement duration in the chronic stage (figure 2 left panel). Importantly, all kinematic measures showed improvement from the sub-acute stages, to chronic stages (figure 2 right panel). These results indicate that functional impairments continue to improve even after the resolution of synergies, though they remain abnormal (Figure 2).

**Figure 2.**
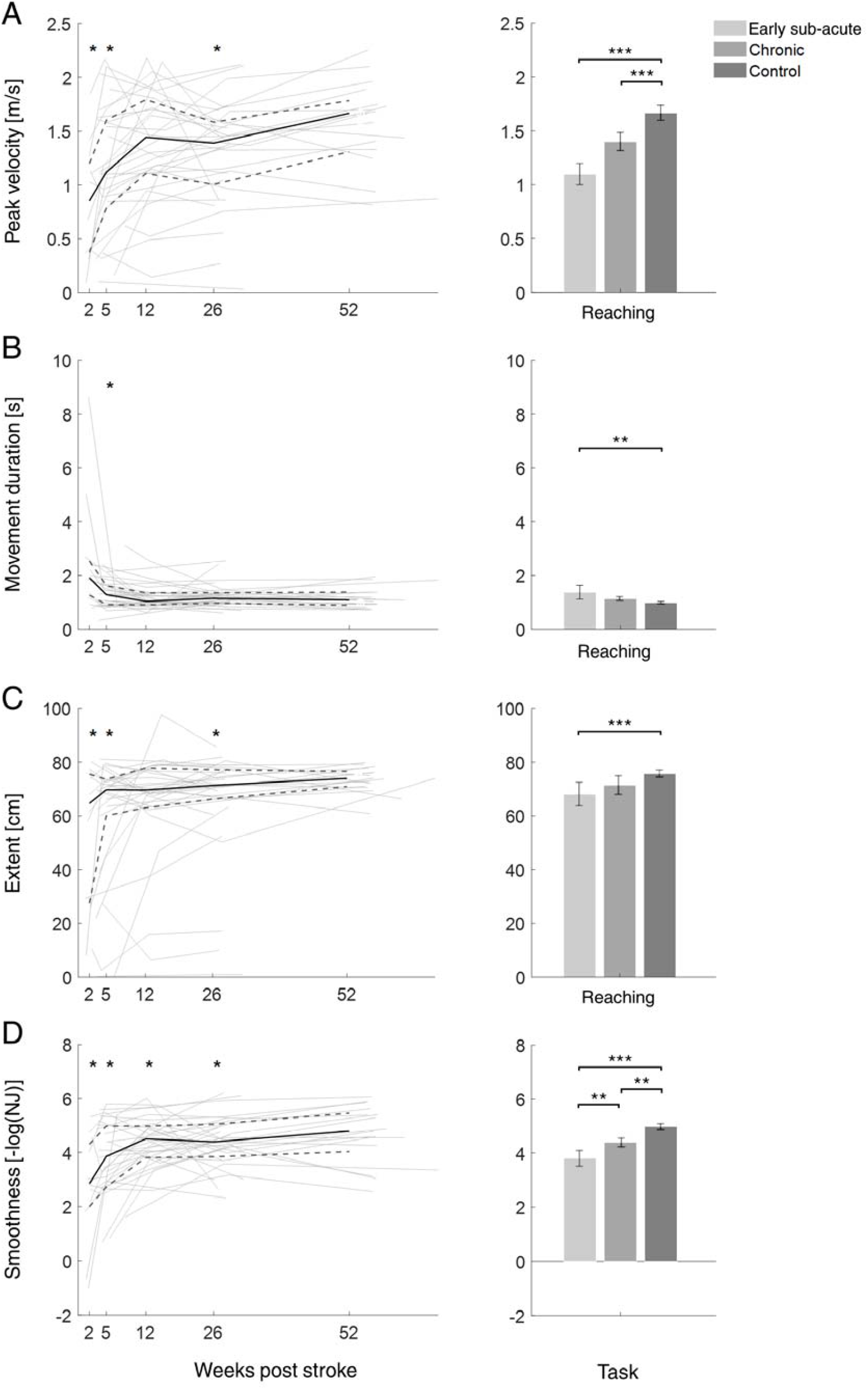
Kinematic measures in different stages of stroke recovery. Plots of (**A-B**) peak velocity, (**C-D**) movement duration, (**E-F**) extent, and (**G-H**) smoothness (minus log of the normalized jerk (NJ)) in the paretic side of stroke participants. Left column: Time course plots for the paretic arm recorded at different stages of recovery. Each line connects the dots of the values of the measure for a single participant across all time points. Group median values are represented by the bold lines, and the group 25^th^ and 75^th^ percentile values are represented by the bottom, and top dashed lines, respectively. Significant one-sample t-tests are denoted by asterisks (* = *P* < 0.01) for each time point (early sub-acute, late sub-acute, and chronic stages). Right column: Group median values of all participants in the early sub-acute stage (1-8 weeks), chronic stage (24-56 weeks), and age-matched controls are denoted in the bar plots. Significant group differences are denoted by asterisks (* = *P* < 0.0167, ** = *P* < 0.01, *** = *P* < 0.001, two-sample t-test with unequal variance).

### Standard measures of weakness and motor function improved from the acute to the chronic stage

If the release from synergies reflects a preliminary stage of recovery, we would expect to see reduction in clinical impairment measures from the early sub-acute phase through to the chronic stage. Most participants exhibited abnormal clinical measures of motor function (ARAT and grip dynamometry) (Figure 3A&C).

**Figure 3.**
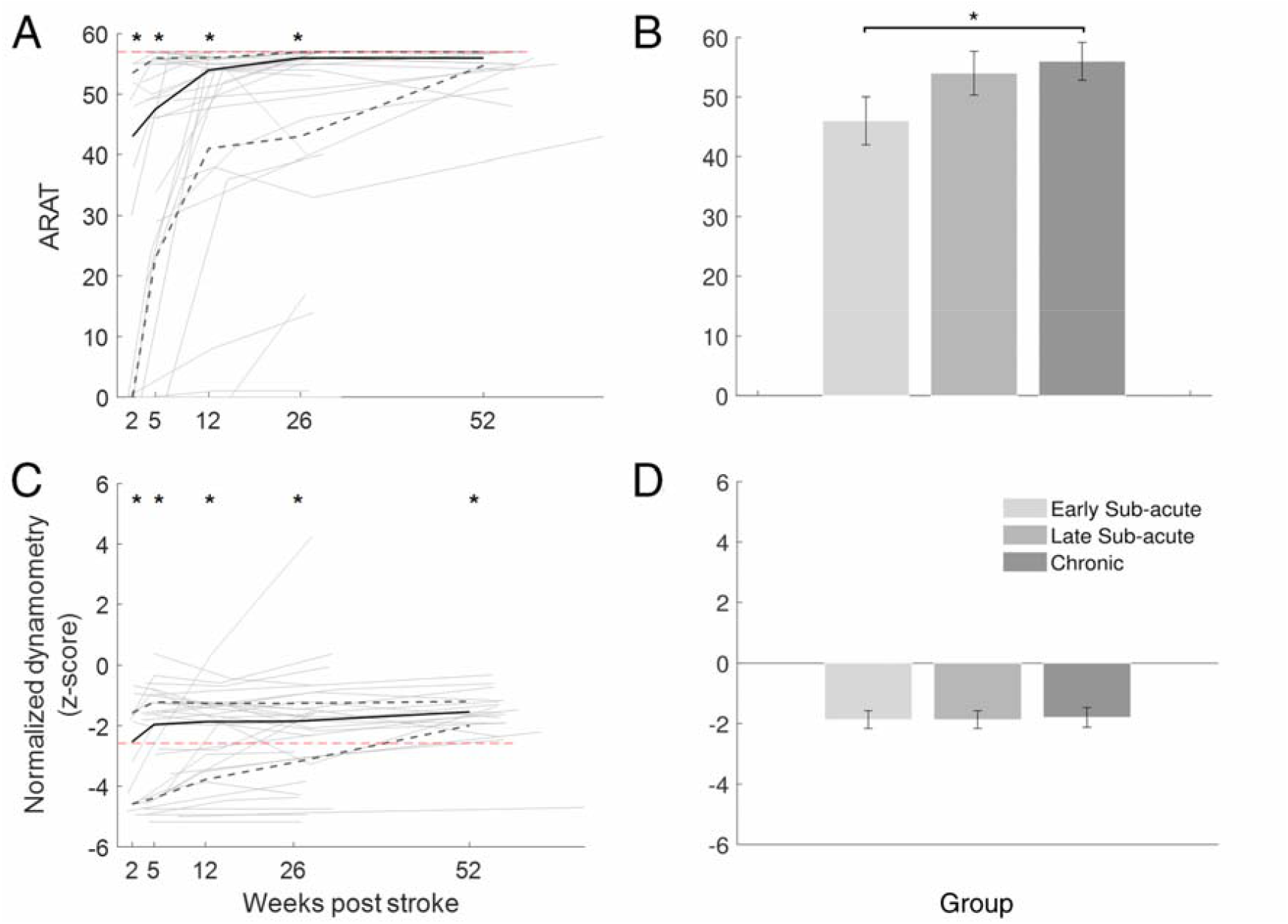
Recovery profiles for strength and functional measures. **(A-B**) ARAT clinical scores time course plots and bar plots. (**C-D)** Grip dynamometry measure time course plots and bar plots. Left column: Time course plots of the paretic arm recorded at different stages of recovery. Each line connects the dots of the values of the measure for a single participant across all time points. Group median values are represented by the bold lines, and the group 25^th^ and 75^th^ percentile values are represented by the bottom, and top dashed lines, respectively. For ARAT scores, the maximum score (57) is denoted in a red dashed line at the top. For normalized dynamometry a threshold a two-tailed test at an alpha level of 0.01 (z-score = -2.58) is denoted in a red dashed line at the bottom. Significant one-sample t-tests are denoted by asterisks (* = *P* < 0.01) for each time point (early sub-acute, late sub-acute, and chronic stages). Right column: Group median values of all participants in the early sub-acute stage (1-8 weeks), late sub-acute stage (11-15 weeks), chronic stage (24-56 weeks), are denoted in the bar plots. Significant group differences are denoted by asterisks (* = *P* < 0.0167, ** = *P* < 0.01, *** = *P* < 0.001, two-sample t-test with unequal variance).

At the early sub-acute stage, 21 of the participants had ARAT scores lower than 50. Most impaired participants improved, but scores were still abnormal in the chronic stage: 14 participants scored the maximum scores for ARAT (57) (Figure 3A). These are consistent with the reported recovery patterns as assessed by ARAT.^19^

Grip dynamometry scores were also abnormal from the early sub-acute stage throughout recovery to later chronic stages (*P* < 0.0001) (Figure 3C).

### The contributions of weakness and flexor synergy intrusion to motor function varied over the time-course of recovery

Next, we examined the association between synergy and weakness recovery on an individual basis. We focused on participants with severe deficits (ARAT below 21)^24^ at the early sub-acute and chronic stages (Figure 4). In the sub-acute stage, all severe participants exhibited weakness (strength lower than four standard deviations below the population mean), and most also exhibited intrusion of synergies (flexor synergy strength higher than zero) (Fig. 4). Looking at the participants with severe impairment in the chronic stage, all of them (5/5) still had significant weakness, but only two participants out of five had flexor synergy intrusion (Fig. 4). This analysis indicates that intrusion of flexion synergy affected motor function in the earlier stages of recovery, whereas weakness was associated with reduced function throughout.

**Figure 4.**
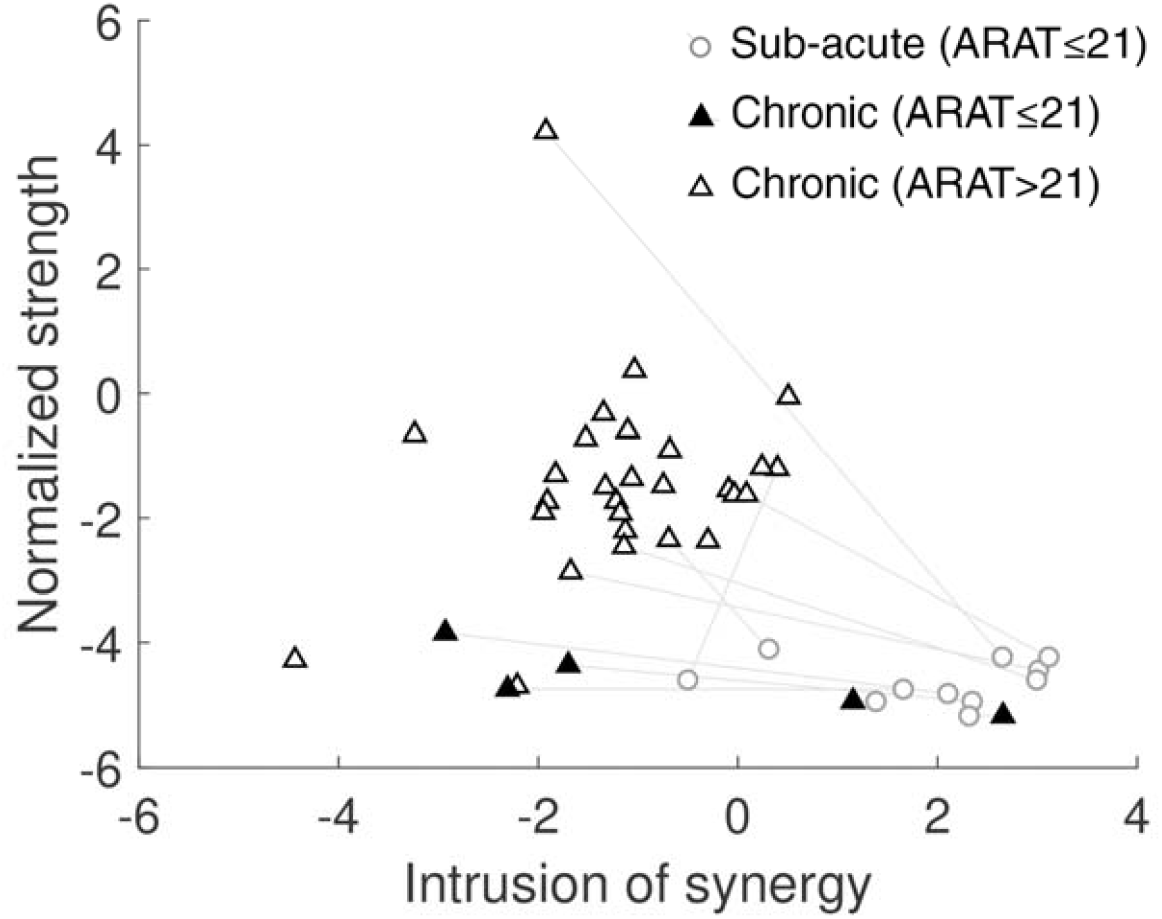
Impaired participants exhibit weakness and intrusion of synergies. Scatter plot of the strength measure compared to the flexor synergy strength measure in the chronic stage (open and closed triangles). Participants with ARAT scores lower than 21 are depicted by closed triangles and with ARAT higher than 22 are depicted by open triangles. Participants with severe impairment in the sub-acute phase (ARAT lower than 21) are presented in faded circles that are connected by a line to their chronic results. Each triangle\circle represents an individual participant.

Finally, to further quantify the differential contribution of weakness and synergies to impairment following stroke, we performed a regression analysis of functional and impairment measures as the dependent variables and weakness and synergies as the two independent variables (Tables 3 & 4). First, we measured the contribution of weakness and synergy to clinical measures. The regression model was significant for both ARAT and Fugl-Meyer scores at all time points (R^2^ > 0.57, *P* < 0.004). Weakness significantly explained both dependent variables at all three time points (*P* < 0.01). However, intrusion of flexor synergy was a significant predictor only for the early and late sub-acute stages but not the chronic stage (Table 3).

**Table 3.** Regression analysis of impairment measures.

| Kinematic measure | R squared | P value | Intercept |  |  | Weakness<br>(dynamometer grip) |  |  | Flexor synergy<br>strength reaching |  |  |
| --- | --- | --- | --- | --- | --- | --- | --- | --- | --- | --- | --- |
|  |  |  | beta | std | p | beta | std | p | beta | std | p |
| ARAT |  |  |  |  |  |  |  |  |  |  |  |
| Early sub-acute | 0.84 | 0.000 | 51.01 | 3.78 | 0.000 | 5.83 | 1.31 | 0.000 | -8.17 | 1.36 | 0.000 |
| Late sub-acute | 0.73 | 0.000 | 51.07 | 4.50 | 0.000 | 4.71 | 1.50 | 0.003 | -8.77 | 1.81 | 0.000 |
| Chronic | 0.58 | 0.003 | 51.40 | 5.83 | 0.000 | 6.96 | 1.56 | 0.000 | -2.31 | 2.21 | 0.305 |
| Fugl-Meyer |  |  |  |  |  |  |  |  |  |  |  |
| Early sub-acute | 0.80 | 0.000 | 65.86 | 4.25 | 0.000 | 7.32 | 1.47 | 0.000 | -6.36 | 1.53 | 0.000 |
| Late sub-acute | 0.79 | 0.000 | 61.87 | 3.30 | 0.000 | 5.04 | 1.10 | 0.000 | -6.58 | 1.33 | 0.000 |
| Chronic | 0.59 | 0.003 | 57.53 | 5.30 | 0.000 | 6.26 | 1.41 | 0.000 | -2.49 | 2.01 | 0.224 |

\*Significant models and independent variables using an alpha value of 0.05 are marked in bold font. Statistics are calculated for the early sub-acute (1 – 8 weeks post-stroke), late sub-acute (11 – 16 weeks post-stroke) and chronic (24 – 56 weeks post-stroke) stages.

Repeating this analysis with peak velocity and extent as the dependent variables produced similar results (Table 4). Weakness significantly explained peak velocity and extent at most sub-acute and chronic time points. Synergies were only significant at the sub-acute stages (Table 4).

**Table 4.** Regression analysis of impairment measures and kinematic measures.

| Kinematic measure | R squared | P value | Intercept |  |  | Weakness<br>(dynamometer grip) |  |  | Flexor synergy strength<br>reaching |  |  |
| --- | --- | --- | --- | --- | --- | --- | --- | --- | --- | --- | --- |
|  |  |  | beta | std | p | beta | std | p | beta | std | p |
| Extent |  |  |  |  |  |  |  |  |  |  |  |
| Early sub-acute | 0.77 | 0.000 | 69.28 | 5.03 | 0.000 | 4.40 | 1.74 | 0.014 | -10.02 | 1.82 | 0.000 |
| Late sub-acute | 0.49 | 0.000 | 69.60 | 6.32 | 0.000 | 3.40 | 2.11 | 0.112 | -7.83 | 2.54 | 0.003 |
| Chronic | 0.45 | 0.021 | 68.13 | 7.52 | 0.000 | 6.11 | 2.01 | 0.005 | -4.46 | 2.85 | 0.127 |
| Peak velocity |  |  |  |  |  |  |  |  |  |  |  |
| Early sub-acute | 0.70 | 0.000 | 1.44 | 0.13 | 0.000 | 0.15 | 0.04 | 0.001 | -0.16 | 0.05 | 0.001 |
| Late sub-acute | 0.31 | 0.014 | 1.68 | 0.20 | 0.000 | 0.15 | 0.07 | 0.033 | -0.08 | 0.08 | 0.325 |
| Chronic | 0.50 | 0.012 | 1.56 | 0.17 | 0.000 | 0.17 | 0.04 | 0.000 | -0.01 | 0.06 | 0.884 |

\*Significant models and independent variables using an alpha value of 0.05 are marked in bold font. Statistics are calculated for the early sub-acute (1 – 8 weeks post-stroke), late sub-acute (11 – 16 weeks post-stroke) and chronic (24 – 56 weeks post-stroke) stages.

## Discussion

Characterizing the time courses of development and resolution of weakness and flexor synergies, as well as their relationship to functional recovery after stroke, has been of significant interest to the scientific community since the earliest attempts to track stroke recovery.^6,25^ We quantified these two components longitudinally in a group of 33 participants using kinematic measures. Our findings indicate that motor synergies emerge and resolve during the early stages of recovery, while strength and function continue to improve into the chronic phase. In Twitchell’s original study, in which he followed the recovery of 25 participants after stroke, all of those who regained voluntary movement initially displayed flexor synergy intrusion. Nevertheless, most of these participants recovered sufficiently to perform isolated joint movements. However, four of them could not progress beyond the flexor synergy, even one and a half years after stroke. Inspired by Twitchell’s seminal study, we kinematically assessed flexor synergy intrusion over time in our participants, as well as evaluating the contribution of these synergies to motor function. Similar to Twitchell’s findings, among participants with severe initial deficits, almost all those who showed good recovery by the chronic stage exhibited flexor synergy intrusion in the sub-acute stage (Figure 4). Two of the severely affected participants who did not recover also had persistent intrusive synergies in the chronic stage (closed triangles in Figure 4), but overall poor recovery in the chronic phase was not associated with flexor synergy intrusion. This is interesting as it suggests that persistence of weakness does not inevitably result in persistence of synergies, with the latter previously conjectured to be the consequence of compensatory reticulospinal tract (RST) upregulation in the setting of corticospinal tract (CST) damage-related weakness.^26,27^ Instead, synergies might be an independent read-out of the degree of residual ipsilesional CST control over spinal cord circuitry.^28^

A central strength of this study is the quantitative assessment of the flexor synergy using naturalistic 3D movements throughout the recovery period. The use of naturalistic 3D tasks is important, considering the variety of operational definitions of flexor synergies and the diversity of methodologies employed to assess them. Our results are consistent with other studies that investigated either finger individuation or independent joint control, indicated by the rate of work area reduction as a function of shoulder abduction loading,^7,29^ showing emergence of synergies in the early sub-acute stage followed by substantial recovery within the first three months.

Our results do not imply that pathological flexor synergies do not exist in the chronic stage of stroke. In fact, abnormal individuation of upper limb joints in the chronic stage is a measure of flexor synergy intrusion and is quite common in the clinic.^9^ As discussed above, two of our participants with severe initial deficits had persistent synergies in the chronic phase. The paucity of synergy intrusion in the chronic stage in our study might be explained by the fact that this is a longitudinal study of participants that received a higher than standard dose rehabilitation – a mean of 53 days of inpatient rehabilitation, which may have improved their chances of overcoming their flexor synergies. In cross-sectional chronic studies, which typically recruit participants with residual motor deficits, there is a bias towards selecting for participants with inferior recovery, which may be associated with persistent flexor synergies. Thus, cross-sectional studies will underestimate the number of chronic stroke participants with good recovery and correspondingly overestimate the persistence of flexor synergy intrusion in this group.

This study offers a novel way to quantify abnormal flexor synergies that could be incorporated into the clinical setting and allow phenotyping of the stroke deficit over the time course of recovery. The dissociation between weakness and synergies suggests that they may need to be targeted separately during rehabilitation. Our data also suggest that higher doses of upper-limb rehabilitation, as our participants received due to their prolonged hospital stay, might favorably change the time course of synergy development.

## Limitations

Longitudinal studies are challenging and prone to missing data points. As demonstrated in Table 2, we were unable to recruit all participants from the two-week time point (n = 12) and could not record all participants at the 1-year mark (n = 17). To address this caveat, we aggregated the data points collected in the early sub-acute (4 weeks), late sub-acute (13 weeks), and chronic (27 weeks) post-stroke stages for the statistical models and group comparisons (see Tables 3-4 and bar graphs in Figures 1 & 2).

In our longitudinal study, it is likely that the participants have received more than a typical dose of neurorehabilitation, which might have reduced the degree of flexor synergy contribution to reduced function in the chronic phase. This however is not really a limitation but a finding; the evolution of flexor synergy expression is not fixed for a given initial deficit.

Our objective was to investigate whether the joint flexor synergy had a detrimental impact on reaching and motor function. For that purpose, kinematic analysis was used, which allowed us to phenotype the emergence of flexor synergy in a naturalistic 3D reaching task over the time-course of recovery. While we posit that our kinematic intrusion measure would be associated with an abnormal EMG signature^30–32^ the converse does not follow: abnormal EMG signals in isometric tasks may not be associated with intrusion of synergies during unconstrained 3D reaching movements.

## Data availability

The data that support the findings of this study are available from the corresponding author, upon reasonable request.

## Acknowledgements

The study was supported by The Lillian and David E. Feldman Research fund.

## Funding

United States - Israel Binational Science Foundation 2021248 to AA and LS

Israel science foundation 1244/22 to LS

